# Frequency-resolved cortical functional connectivity across the adult lifespan

**DOI:** 10.1101/2025.03.28.645908

**Authors:** Santeri Ruuskanen, Juan Camilo Avendano-Diaz, Mia Liljeström, Lauri Parkkonen

## Abstract

The operation of the human brain relies on functional networks enabled by inter-areal oscillatory synchronization between neuronal populations. Although disruptions in this functional connectivity are associated with brain disorders, evidence on its healthy age-dependent variation and behavioral relevance remains limited. Utilizing magnetoencephalography (MEG) recordings from 576 adults aged 18–87 years, we investigated the evolution of resting-state functional connectivity (rs-FC) across the healthy adult lifespan. We observed age-related, frequency-specific changes in widespread cortical networks. Alpha-band (8–13 Hz) rs-FC decreased, while delta (1–4 Hz), theta (4–8 Hz), and gamma-band (40–90 Hz) rs-FC increased with age. Beta-band (13–30 Hz) rs-FC followed a non-linear trajectory, peaking in middle age. The global delta, theta, alpha, and beta-band patterns differed from concurrent changes in oscillatory power, underscoring their dissociable contributions. Notably, reduced beta-band rs-FC was associated with increased sensorimotor attenuation, indicating that changes in rs-FC are behaviorally relevant for sensorimotor function. These findings advance our understanding of healthy brain aging and highlight a link between resting-state brain activity and sensorimotor integration.

**Key points:**

- Functional connectivity is altered across the healthy adult lifespan in a frequency-dependent manner
- Changes in source power do not explain global functional connectivity trajectories
- Beta-band connectivity at rest is associated with sensorimotor attenuation independent of age-related effects

## Introduction

The healthy aging human brain undergoes widespread structural and functional changes which are further modified in various neuropathologies (Bethlehem et al., 2022; Betzel et al., 2014; Damoiseaux et al., 2008; Ferreira & Busatto, 2013; Mandal et al., 2018; Rempe et al., 2023; Stam, 2010; Stam et al., 2006; Stoffers et al., 2008; Vlahou et al., 2014). It is now well established, based on magnetic resonance imaging (MRI), that cortical thickness and gray matter volume decrease throughout the adult lifespan (Bethlehem et al., 2022). Moreover, diffusion-weighted MRI and functional MRI (fMRI) studies have revealed age-related trends in structural and functional brain networks (Betzel et al., 2014; Damoiseaux et al., 2008; Ferreira & Busatto, 2013). In addition, evidence from non-invasive electrophysiological techniques including electroencephalography (EEG) and magnetoencephalography (MEG) suggests that spontaneous neuronal oscillations and aperiodic activity possess unique frequency-dependent trajectories across the lifespan (Gómez et al., 2013; Rempe et al., 2023; Scally et al., 2018; Thuwal et al., 2021; Vakorin et al., 2025; Vlahou et al., 2014),. However, research on age-related trends in the inter-areal functional coupling between these spontaneous oscillations, referred to as resting-state functional connectivity (rs-FC), and their behavioral relevance remains limited. Nevertheless, functional connectivity between brain areas is vital for normal brain function and disrupted in neuropathological conditions typically associated with aging, such as Parkinson’s and Alzheimer’s disease (Buzi et al., 2023; Mandal et al., 2018; Stam, 2010; Stam et al., 2006; Stoffers et al., 2008). Accurately characterizing the healthy variability in rs-FC is essential to distinguish normal aging from pathological deviations and to optimally utilize rs-FC as a potential disease biomarker.

Changes in MEG rs-FC related to healthy aging have been sparsely investigated, and the findings remain inconclusive. An early study reported age-related increases in directed rs-FC into the medial temporal lobe and decreased rs-FC into the posterior cingulum and precuneus across broad frequency ranges within 1–100 Hz (Schlee et al., 2012a). Others have reported decreased sensor-level global coherence in the alpha frequency band across the lifespan (Sahoo et al., 2020) and increased variability of the phase-locking value (PLV) within the default mode network (DMN) in the delta frequency band in older adults (Jauny et al., 2022a). In contrast, a more recent study found no significant age-related changes in the imaginary part of coherency (ImC) in the delta band, and reported increased connectivity of posterior, parietal, and temporal regions in the theta and gamma bands, as well as decreased alpha and beta connectivity with age in temporal and posterior areas, respectively (Stier et al., 2023). Another recent report suggested more complex patterns of generally decreasing connectivity in the delta band, nonlinear trends in the theta and alpha bands, and increasing connectivity in the beta and gamma bands (Doval et al., 2024). Notably, in contrast to these studies utilizing connectivity metrics based on phase synchronization, Coquelet and colleagues (2017) did not observe any significant age-related changes in functional connectivity using power envelope correlations. In addition to MEG, age-related changes in rs-FC have been reported in EEG (Ishii et al., 2018). However, while decreased alpha-band connectivity in the elderly has also been observed in EEG (Moezzi et al., 2019; Scally et al., 2018), effects in other frequency bands are inconsistent across studies and in comparison to the MEG literature (Moezzi et al., 2019; Samogin et al., 2022).

Despite common elements across the findings of previous MEG rs-FC studies, research in the area has not yet converged. This incongruity could be largely attributed to methodological differences between studies. Besides limited sample sizes, we identified four main methodological considerations that could confound rs-FC studies. First, a major challenge in electrophysiological functional connectivity analysis stems from source leakage, which may lead to false positive findings in the absence of true synchronization (Palva et al., 2018). Several groups have utilized functional connectivity estimation methods highly susceptible to false positive findings due to source leakage, such as coherence and PLV. Second, some studies have been limited to sensor-level analyses, which accentuates the linear mixing between signals leading to spurious connectivity estimates and compromises the neurophysiological interpretability of the results (Schoffelen & Gross, 2009). Third, many previous studies have used coarse anatomical parcellations, which results in signal cancellation as source time courses are averaged within parcels. Finally, results on rs-FC could be confounded by concurrent changes in oscillatory source power, which affects functional connectivity estimates through altered signal-to-noise ratio (Donoghue et al., 2022).

The presence of putative age-related alterations in functional connectivity elicits interest in the potential connection between rs-FC and the maintenance of cognitive and sensorimotor function in older age. Age-related increases in functional connectivity during task performance and rest have generally been associated with the maintenance of cognitive performance, while decreased connectivity has been observed in patients with cognitive decline (Buzi et al., 2023; Jauny et al., 2022b; Mandal et al., 2018). Nevertheless, the magnitude and direction of the effects are typically frequency-band specific and vary between studies. Most studies investigating age-related changes in rs-FC also report associations between cognitive performance and connectivity. In particular, reduced visual short-term memory (VSTM) performance has been related to decreased alpha-band global coherence and increased delta-band PLV variability in bilateral supramarginal regions (Jauny et al., 2022a; Sahoo et al., 2020). On the other hand, weaker performance in tasks testing fluid intelligence and visual memory has been reported with increased bilateral temporal lobe connectivity in the beta and gamma frequency ranges and reduced directed connectivity into posterior medial areas in the 1–100-Hz frequency range (Schlee et al., 2012a; Schlee et al., 2012b). Others have also reported associations between the similarity of structural and functional networks and higher performance in the mini-mental state evaluation (MMSE; Folstein et al., 1975) and VSTM tasks, suggesting that similar age-related reorganization of structural and functional connectivity has cognitive benefits (Jauny et al., 2024).

Aging is typically associated with reduced sensory sensitivity (Konczak et al., 2012). The combination of perceived sensory stimuli and internal forward models of movement is known as sensorimotor integration (Wolpert et al., 1995), where the contributions of sensory feedback and prediction are weighted by their relative precision (Körding & Wolpert, 2004). As a result, self-generated actions are perceived as weaker than externally generated actions (Chapman et al., 1987). This phenomenon is known as sensorimotor attenuation. Reduced sensory sensitivity with age leads to increased reliance on prediction and, hence, to increased sensorimotor attenuation (Wolpe et al., 2016). Anatomical and functional MRI data indicate that this effect is associated with decreased gray matter volume in the supplementary motor area (pre-SMA) and decreased fMRI functional connectivity between the pre-SMA and frontostriatal regions during both resting state and a movement task (Wolpe et al., 2016). However, despite the prominent role of beta oscillations in sensorimotor control (Barone & Rossiter, 2021; Engel & Fries, 2010), the electrophysiological underpinnings of the age-related increase in sensorimotor attenuation remain elusive.

The aims of the current study were twofold. First, we aimed to elucidate the trends of rs-FC across the healthy adult lifespan. Specifically, we employed MEG to investigate the trajectories of source-space rs-FC across the healthy adult lifespan in the Cam-CAN (Cambridge Centre for Ageing and Neuroscience) cohort (Shafto et al., 2014; Taylor et al., 2017). We analyzed resting-state MEG recordings from a cross-sectional sample of 576 participants aged 18–87 years from the Cam-CAN database (**Fig 1A**). We addressed the common methodological limitations including source leakage, suboptimal cortical parcellation, and confounding oscillatory power. To this end, we estimated neuronal source time courses from preprocessed MEG data using the dynamic statistical parametric mapping (dSPM) inverse method. We then estimated rs-FC between all pairs of 448 subregions using weighted phase-lag index (WPLI) as the connectivity metric in the delta (1–4 Hz), theta (4–8 Hz), alpha (8–13 Hz), beta (13–30 Hz), and gamma (40–90 Hz) frequency bands (**Fig 1B**). We examined the putative linear and nonlinear relationships between age and rs-FC using linear and quadratic association measures based on hierarchical regression. Additionally, we analyzed age-related trends in neural source power to determine whether they mirrored those of rs-FC, which would hinder the interpretation of changing rs-FC patterns as true changes of inter-areal connectivity.

**Figure 1.**
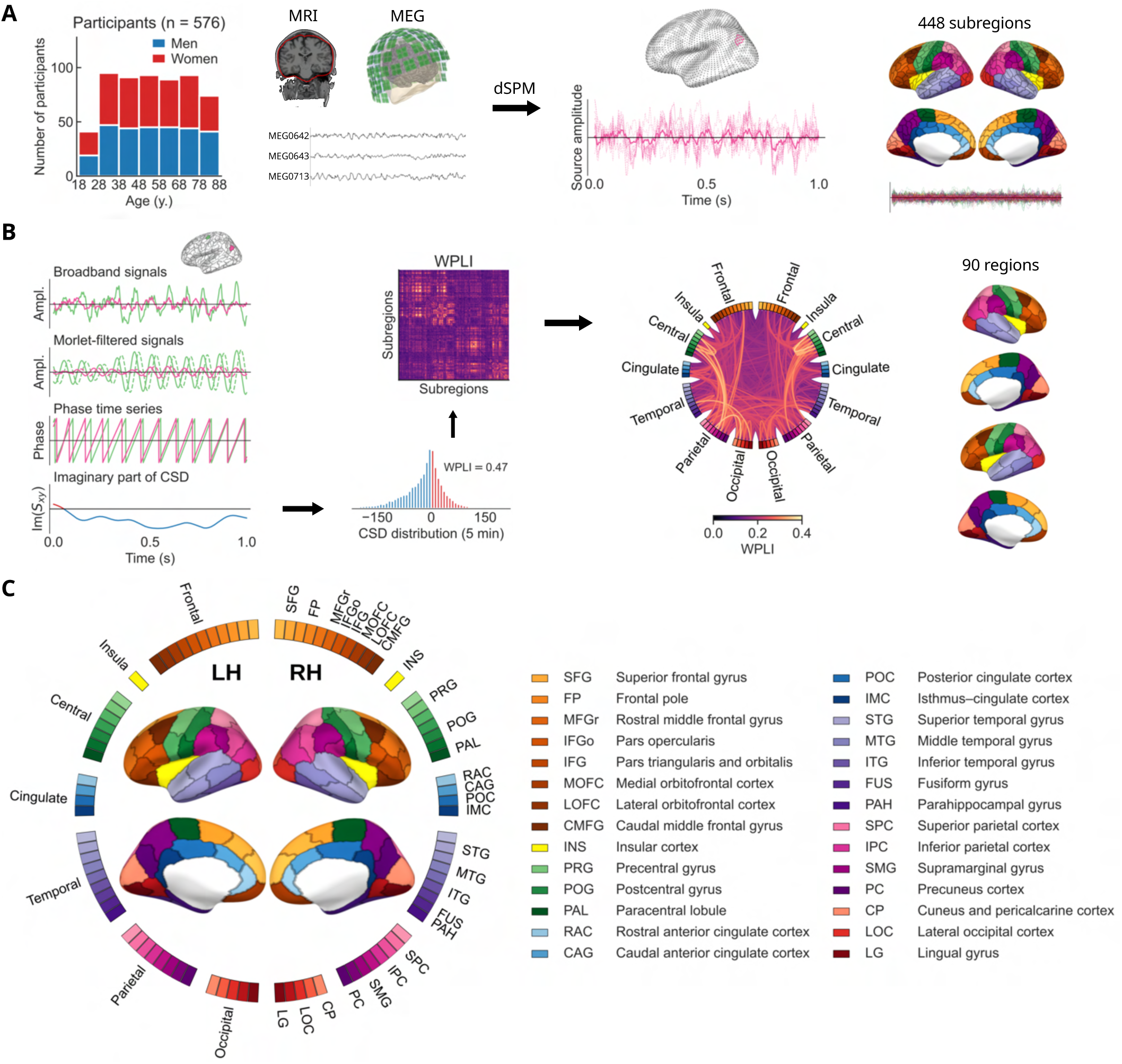
Data and analysis pipeline. (**A**) Resting-state MEG and structural MRI data from 576 healthy participants were used to extract neuronal source time courses at 5124 cortical source points. The source time courses were spatially averaged within 448 cortical subregions based on an anatomical parcellation. (**B**) Analytic signals in the 1–100-Hz frequency range were obtained using Morlet wavelet filtering, and the cross-spectral density (CSD) was calculated for each pair of subregions. The imaginary part of CSD was used to calculate the Weighted Phase-Lag Index (WPLI) across time, resulting in a 448 × 448 all-to-all connectivity matrix where each link quantifies the asymmetry of the phase difference distribution across time. The WPLI values were spatially averaged within a sparser parcellation of 90 cortical regions, suitable for presentation in a circular graph. (**C**) Parcellation reference. Cortical regions are placed around the circle and the lines between them represent connections in the network.

Our second aim was to explore the potential role of rs-FC in the increased sensorimotor attenuation observed in older age (Wolpe et al., 2016). To this end, we analyzed the relationship between rs-FC and force overcompensation in a force matching task, which is a well-stablished measure of sensorimotor attenuation. Given that beta-band connectivity both at rest and during task performance has been associated with accurate motor performance (Chung et al., 2017; Samogin et al., 2022), we hypothesized that the baseline rs-FC levels could be indicative of performance in the force matching task and reflect the efficient integration of sensory signals.

## Materials and Methods

### Participants and recordings

The data were collected at the Medical Research Council (UK) Cognition and Brain Sciences Unit (MRC-CBSU) in Cambridge, UK, as part of the Cambridge Center for Ageing and Neuroscience (Cam-CAN) study (Shafto et al., 2014; Taylor et al., 2017). The cross-sectional, multimodal open-access database includes structural and functional MRI, MEG, and behavioral recordings of a population-based cohort of nearly 700 healthy participants aged between 18 and 88 years with an approximately uniform age distribution. The participants were selected for neuroimaging and detailed behavioral experiments following an initial screening stage that assessed general and cognitive health. The inclusion criteria comprised normal cognitive health, no serious psychiatric conditions, meeting hearing and English language requirements, and no contraindications for MRI recordings (for details, see (Shafto et al., 2014)). In our primary analysis, we included *n* = 576 participants after excluding participants due to missing anatomical MRI or resting-state MEG data, poor data quality, missing head position indicator (HPI) signals or electrooculogram (EOG) channels, and failed FreeSurfer reconstructions (**Fig 1A**). To investigate the relationship between functional connectivity and sensorimotor attenuation, we included a subset of *n* = 284 participants with available recordings from the force matching task collected in the Cam-CAN study. The primary study was conducted in accordance with the Declaration of Helsinki and was approved by the Cambridgeshire 2 Research Ethics Committee (reference: 10/H0308/50). The participants gave written informed consent.

MRI data were collected at MRC-CBSU using a 3-T Siemens TIM Trio scanner (Siemens Healthineers AG, Munich, Germany) and a 32-channel head coil. T1-weighted scans were collected using a magnetization-prepared gradient echo sequence with the following parameters: repetition time 2250 ms, echo time 2.99 ms, flip angle 9^◦^, and inversion time 900 ms. MEG data were recorded in the seated position during rest (eyes closed) using a 306-channel VectorView MEG system (Megin Oy, Espoo, Finland) at MRC-CBSU. Horizontal and vertical EOG and single-lead electrocardiogram (ECG) were recorded simultaneously with MEG. The head position of the participant was continuously monitored using four HPI coils. The duration of the resting-state recording was 8 minutes and 40 seconds. Data were bandpass-filtered to 0.03–330 Hz and sampled at 1 kHz. Further details on the MRI and MEG data acquisition are available in the publication by Taylor and colleagues (2017).

The force matching task, which quantifies sensorimotor attenuation, was performed in a separate session from the neuroimaging measurements by half of the neuroimaging participants (Shafto et al., 2014). In the experimental setting, a torque motor attached to a lever was used to apply a target force to the participant’s left index finger, which rested below the lever. Following concurrent auditory and visual cues, the participant attempted to match the target force with their right index finger either by directly pressing on the lever (direct condition) or by operating a linear potentiometer connected to the torque motor (indirect condition). A force sensor attached to the lever recorded both the target and matched forces. The experiment included 32 trials for each condition. The difference between the target and matched forces is referred to as overcompensation. An absence of overcompensation would indicate that the participant matched the target force perfectly. Positive values in the direct condition indicate sensorimotor attenuation, while the indirect condition serves as a control for the sensitivity of haptic pressure perception (Wolpe et al., 2016). In our analysis, we only considered the direct condition as a measure of the extent of sensorimotor attenuation.

### MEG preprocessing

We followed standard MEG analysis practices, consisting of data preprocessing, coregistration with structural MRI, and estimation of neural current sources in a source space based on individual brain anatomy. The analysis was performed primarily using MNE-Python software version 1.6 (Gramfort et al., 2013; Gramfort et al., 2014).

Spatiotemporal signal-space separation (tSSS; Taulu & Simola, 2006) was employed to remove external interference and compensate for head movement during the measurement (MaxFilter v2.3 (Megin Oy, Espoo, Finland); correlation limit 0.98; correlation window length 10 s). Data were bandpass-filtered to 1–100 Hz and notch-filtered at the 50-Hz line frequency and harmonics. The same procedure was applied to the resting-state and empty-room data, except for head movement correction. Independent Component Analysis (ICA) was used to dampen artifacts arising from eye movement and cardiac activity (FastICA; Hyvärinen, 1999). Independent components were automatically selected for removal based on their Pearson correlation with the EOG channels and cross-trial phase statistics with the ECG channel (Dammers et al., 2008). Up to two components corresponding to each artifact modality were removed. Following artifact removal, the MEG data were divided into 30-s epochs, and any epochs containing residual artifacts were rejected based on peak-to-peak amplitude thresholds (magnetometers: 10 pT; gradiometers: 10 pT/cm). The first 10 good 30-s epochs for each subject were included in the subsequent analysis for a total recording duration of 5 min. Subjects with less than 10 good epochs were excluded from further analysis, resulting in a sample size of *n* = 576.

### MRI processing and forward model

T1-weighted structural MRI data were used to create 3-D cortical reconstructions of each participant using FreeSurfer software (Dale et al., 1999; Fischl et al., 1999). For the MEG forward model, we used a single-compartment linear collocation boundary element method (BEM) volume conductor model. The intracranial volume boundary tessellation was obtained through the FreeSurfer watershed algorithm (Ségonne et al., 2004). A cortically-constrained source space with recursively subdivided icosahedron spacing (ico4) was defined for each participant, resulting in 5124 cortical source points. The anatomical data were manually coregistered with the MEG measurements using the MNE-Python coregistration GUI. The MEG–MRI coregistrations were provided by Bardouille and colleagues (2019).

### Source estimation

We estimated the time courses of neuronal activation using the dynamic statistical parametric mapping (dSPM; Dale et al., 2000) inverse method (regularization parameter 0.111; depth-weighting exponent 0.8; loose orientation constraint 0.2 (Lin et al., 2006)). We used the sample covariance method to estimate noise covariance matrices from empty-room recordings measured in the same session with the recording of each participant. Source time courses were estimated at the 5124 cortical source points and morphed into the FreeSurfer *fsaverage* reference brain template to facilitate analysis across subjects. Both magnetometers and gradiometers were used in source estimation.

### Cortical parcellation

To decrease computational complexity and improve interpretability, we employed a cortical parcellation to reduce the dimensionality of the source-space data. To that end, the data were organized into 448 subregions as defined in the *aparc sub* parcellation scheme developed by Khan and colleagues (2018) based on the Desikan–Killiany (DK) atlas (Desikan et al., 2006) (**Fig 1A**). The time courses of the source points within each subregion were then averaged to form a representative time series. To prevent signal cancellation at opposite sides of sulci, a sign-flip operation was applied to the signals of the sources whose orientation differed from the dominant direction within the subregion by more than 90^◦^. The use of a relatively dense parcellation further reduces signal cancellation and flattening during the averaging procedure.

For the group-level statistical analysis, we used a custom parcellation, which was created by grouping the subregions from the *aparc sub* parcellation (Khan et al., 2018) in a way that avoids elongated regions, thereby maintaining a homogeneous region size across the brain and facilitating interpretable visualizations. This process resulted in a parcellation with 90 regions. The custom parcellation has an intermediate granularity between the *aparc sub* parcellation and the original DK atlas, which improves the anatomical interpretability of the involved cortical areas and limits the overall number of connections in the subsequent statistical analysis. The custom parcellation is illustrated in **Fig 1C**. The names of the resulting regions were inherited from the DK atlas. For regions corresponding to subdivisions of a DK atlas parcel, the names were appended with initial characters of the alphabet.

### Morlet wavelet filtering

Pairwise phase-based functional connectivity metrics, such as coherence and phase-lag index, can be computed from the cross-spectral density between two analytic signals. A narrowband analytic signal represents the activation of a brain region across time in a narrow frequency bin. To obtain the analytic signals, we applied Morlet wavelet filtering to the representative time series of the 448 subregions (**Fig 1B**). The representative time series were convolved with complex Morlet wavelets with the Morlet parameter (number of cycles) *ω* = 5. The fixed-cycle-count Morlet filter is consistent with the phenomenological organization of neuronal oscillations in bursts of a few cycles (Van Ede et al., 2018). The frequency response of the Morlet filter is Gaussian, and its frequency-domain standard deviation scales with the center frequency. We used 32 logarithmically-spaced center frequencies between 1–100 Hz to avoid capturing redundant information while also reducing the computational cost compared to a linearly-spaced frequency grid. The resulting narrowband analytic signals were used to calculate the source power of each subregion and the cross-spectral density between all pairs of subregions. The cross-spectral density was subsequently employed to estimate functional connectivity using the Weighted Phase-Lag Index (WPLI).

### Functional connectivity analysis

Inter-areal synchronization between neuronal oscillations provides a mechanism for functional connectivity between neuronal populations (Fries, 2005; Womelsdorf et al., 2007). The extent of functional connectivity can be quantified by evaluating the consistency of the phase difference between two oscillatory signals representing spatially separated brain areas. We estimated all-to-all pairwise functional connectivity between the 448 subregions using the Weighted Phase-Lag Index (WPLI) as the connectivity metric (Vinck et al., 2011). WPLI quantifies the asymmetry of the phase-difference distribution between a pair of narrowband analytic signals *x*(*f, t*) and *y*(*f, t*) over *N_t_* time points at frequency *f* as

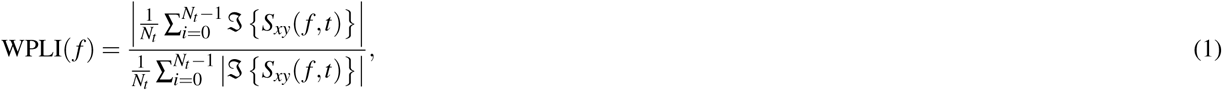

where ℑ{·} denotes the imaginary part and *S_xy_*(*f, t*) is the cross-spectral density *x*(*f, t*)*y*(*f, t*)^∗^ between the signals. The WPLI method minimizes both the influence of artificial zero-lag interactions due to source leakage and the sensitivity to noisy small phase differences (Vinck et al., 2011). WPLI has been widely applied and produces reliable group-level results (Colclough et al., 2016). We applied the implementation of MNE-Connectivity software version 0.5 (https://mne.tools/mne-connectivity/).

We computed WPLI separately in each frequency bin and averaged the results across frequencies within the canonical delta (1–4 Hz), theta (4–8 Hz), alpha (8–13 Hz), beta (13–30 Hz), and gamma (40–90 Hz) frequency bands. To reduce the dimensionality of the 448 × 448 connectivity matrices, we employed a sparser custom parcellation of 90 regions in the region- and connection-level analyses (**Fig 1B**). The 448 × 448 connectivity matrices were transformed into the sparser parcellation by spatially averaging the connectivity scores within each of the 90 regions, yielding 90 × 90 matrices. These matrices constitute the adjacency matrices of the weighted connectivity networks (**Fig 1B**). We analyzed unthresholded networks to avoid the artificial selection of a threshold value. In addition to analyzing individual connections, we considered the mean connectivity of each region, which we defined as the mean strength of the connections associated with the region. Furthermore, we analyzed age-related changes in global connectivity, defined as the mean of all connections in the connectivity network.

Although functional connectivity metrics based on phase synchronization are widely applied, numerous studies have also employed power or amplitude envelope correlations (AEC), including in healthy aging and neurodegenerative disorders (Coquelet et al., 2017; Hipp et al., 2012; Schoonhoven et al., 2022). AEC has also been suggested to have higher test–retest reliability than most phase-based metrics (Colclough et al., 2016; Garcés et al., 2016; Marquetand et al., 2019) and could capture information complementary to phase synchronization (Siems & Siegel, 2020). Therefore, in addition to the primary WPLI analysis, we repeated the functional connectivity analysis using AEC on the same narrowband analytic signals used to estimate WPLI. We calculated AEC between pairwise-orthogonalized amplitude envelopes to avoid artificial zero-lag interactions (Hipp et al., 2012). The results are discussed in Supplementary Material.

### Permutation statistics of linear and quadratic associations

We investigated the trajectories of functional connectivity across the adult lifespan at three levels: individual connections in the connectivity network, mean connectivity of each region, and global connectivity across the whole network. To visually inspect age-related trends, we averaged the connectivity matrices and global connectivity across subjects within each age decade. Given that age-related phenomena are often nonlinear, we assessed both linear and quadratic associations between participant age, functional connectivity, source power, and overcompensation in the force matching task.

We modeled linear and quadratic associations using hierarchical linear regression. First, we fit a simple linear regression model with age as the independent variable to quantify linear associations. At the second level, we considered quadratic associations after removing the linear trend, adding age squared as an independent variable to the model. We evaluated the statistical significance of the associations using a nonparametric two-tailed permutation test. We indexed the effect size of the linear associations using the Pearson correlation coefficient and that of the quadratic associations using the model coefficient of determination multiplied by the sign of the quadratic term coefficient, which were used as test statistics in the permutation test. The permutation distribution was obtained by shuffling the dependent variable vectors, while the independent variables were kept fixed (*n*_permutations_ = 1000). This is equivalent to the null hypothesis that the model does not explain any variance in the data (i.e., the coefficient of determination is zero).

We leveraged a mass univariate approach to analyze the associations between participant age and functional connectivity at the level of individual connections in the connectivity network, fitting a separate hierarchical regression model for each connection. The number of subjects was *n* = 576. This resulted in 3960 fitted hierarchical regression models per frequency band, one for each link in the network. For mean connectivity, a similar approach resulted in 90 fitted models per frequency band. In addition to evaluating the association between participant age and local connectivity, we also assessed the relationship between overcompensation in the force matching task and mean connectivity (*n* = 284). At the whole network level, we fit separate models for each log-spaced frequency bin to provide an association spectrum across the entire 1–100-Hz frequency range, resulting in 32 fitted models. In addition to examining the dependency between age and global connectivity, we also analyzed the relationship between age and mean source power. We corrected all results for multiple comparisons by applying false discovery rate correction (FDR; Benjamini & Hochberg, 1995) separately at each analysis level. At the connection and region levels, *p*-values were pooled across connections and regions within frequency bands, whereas at the whole network level, they were pooled across frequency bins.

### Standardized multiple linear regression analysis of sensorimotor and age effects

Since both overcompensation in the force matching task and rs-FC are likely dependent on participant age, we applied standardized multiple linear regression at the region level to distinguish between sensorimotor and age-related effects on rs-FC. We fit a model of the form *mean connectivity* ∼ *age*, *mean force overcompensation*. The variables were *z*-scored before fitting the model and effect sizes were quantified using the resulting standardized *β* -coefficients. The results were FDR-corrected for multiple comparisons across labels, pooling together the *p*-values pertaining to both explanatory variables. We checked age and mean force overcompensation for collinearity, i.e., that the correlation between them was below 0.7 (Dormann et al., 2013). The analysis was performed using statsmodels version 0.14.5 (Seabold & Perktold, 2010).

## Results

### Functional connectivity across the adult lifespan

We investigated age-related changes in rs-FC at the level of individual network connections. For visual inspection of rs-FC networks across the adult lifespan, we averaged the estimated connectivity matrices across subjects within age cohorts in the delta (1–4 Hz), theta (4–8 Hz), alpha (8–13 Hz), beta (13–30 Hz), and gamma (40–90 Hz) frequency bands (**Fig 2B**). The averaged results showed distinct connectivity patterns in each frequency band. In the delta band, the rs-FC networks were predominantly frontal–temporal. Theta-band networks involved temporal, parietal, and occipital areas. Alpha-band connectivity was most prominent between posterior, parietal, and temporal regions. Notable beta-band rs-FC was observed between centroparietal and occipital areas. Gamma-band connectivity was observed between frontal, temporal, parietal, and occipital areas.

**Figure 2.**
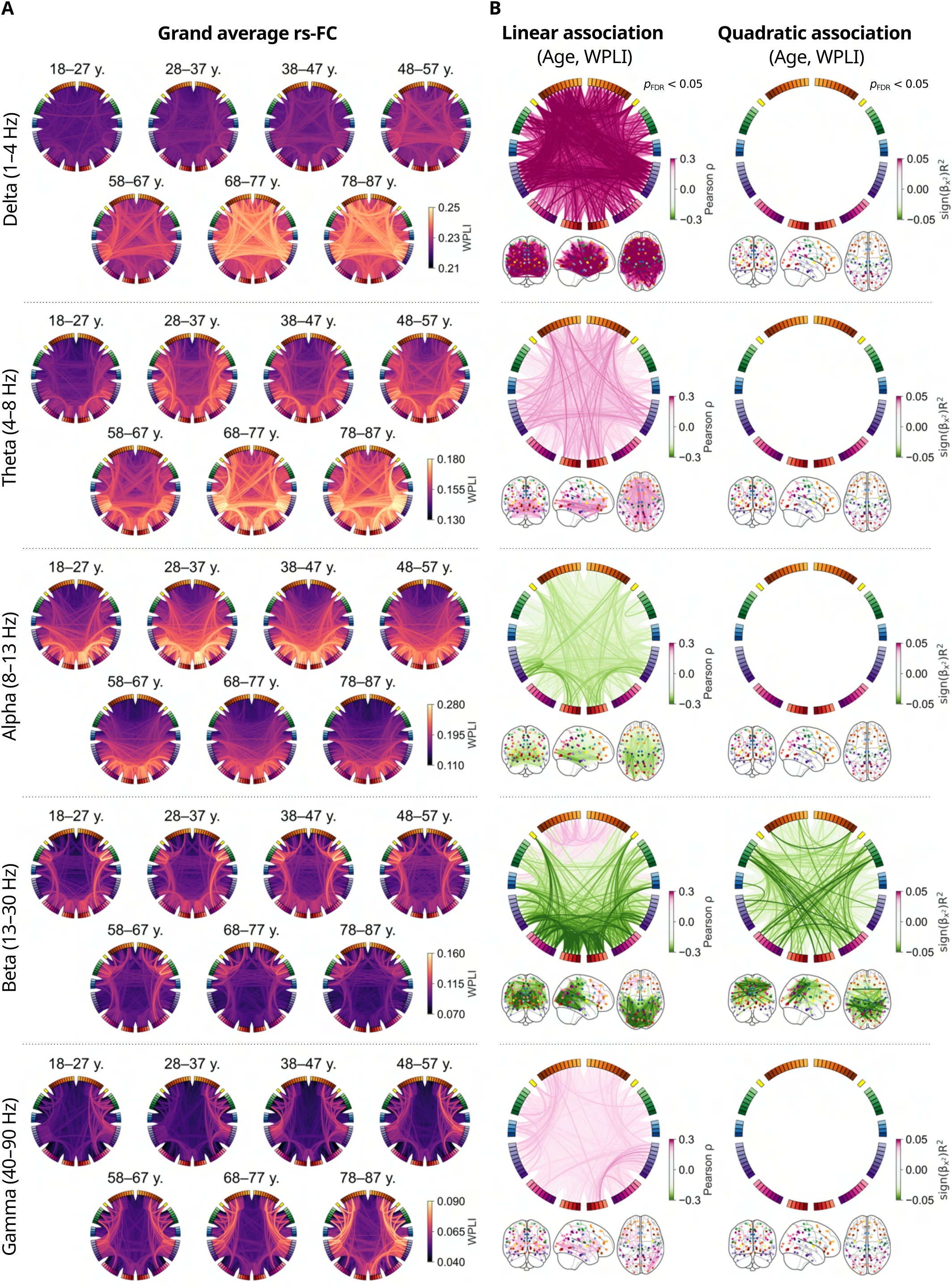
Functional connectivity changes across the adult lifespan. (**A**) Group averages of functional connectivity within age cohorts in the delta (1–4 Hz), theta (4–8 Hz), alpha (8–13 Hz), beta (13–30 Hz), and gamma (40–90 Hz) frequency bands show distinct frequency-dependent patterns. (**B**) Linear (left column) and quadratic (right column) associations between participant age and connections strength reveal frequency-band-specific patterns of change across the adult lifespan. Each colored line in the figure represents the effect size of a significant association; the Pearson correlation coefficient for linear associations, and the coefficient of determination multiplied by the sign of the quadratic term coefficient for the quadratic associations. The circular plots and schematic brains provide complementary visual perspectives of the same data.

To quantify age-related changes in rs-FC across the adult lifespan, we calculated linear and quadratic associations between participant age and the strength of each connection in the rs-FC network in the delta, theta, alpha, beta, and gamma frequency bands (**Fig 2C**). In the delta band, we observed generally increased connectivity. In general, the strongest associations were observed between frontal and temporal areas. The largest effect sizes were found between left isula and left fusiform gyrus (*ρ* = 0.35, *p*_FDR_ = 0.001) and left insula and right middle temporal gyrus (*ρ* = 0.34, *p*_FDR_ = 0.001). The linear effect sizes were generally larger in the delta band compared to other frequency bands. No significant quadratic associations between participant age and rs-FC were observed in the delta band.

In the theta band, we observed increased connectivity, particularly between frontal–temporal and frontal–occipital areas. Notably, a large subset of the connections strengthening with age were interhemispheric. The largest effect size was observed between the left medial orbitofrontal gyrus and the region consisting of the left cuneus and pericalcarine cortex (*ρ* = 0.26, *p*_FDR_ = 0.004). We found similar effects between frontal–temporal areas, including between the left frontal pole and the right middle temporal gyrus (*ρ* = 0.25, *p*_FDR_ = 0.004). No significant quadratic associations were observed between the age of the participants and rs-FC in the theta band.

In the alpha band, we identified linear age-related decreases in rs-FC, most prominently between temporal–occipital areas. Some of the connections associated with the greatest effects were left fusiform gyrus b – left parahippocampal gyrus (*ρ* = −0.27, *p*_FDR_ = 0.004), left lateral occipital b – left parahippocampal gyrus (*ρ* = −0.26, *p*_FDR_ = 0.004), and left lateral occipital a – left fusiform a (*ρ* = −0.26, *p*_FDR_ = 0.004). The letters a and b here refer to the subdivision of Desikan–Killiany atlas regions. We did not observe significant quadratic associations between participant age and rs-FC in the alpha band.

In the beta-band, a general linear decrease in rs-FC was identified, with the strongest effects between posterior and parietal regions. However, contrary to the alpha and theta bands, increased connectivity was observed in the frontal lobe. Some of the connections exhibiting the strongest positive associations with age were left superior frontal gyrus b – left rostral middle frontal gyrus b (*ρ* = 0.22, *p*_FDR_ = 0.003), and left lateral orbitofrontal gyrus – left rostral middle frontal gyrus b (*ρ* = 0.22, *p*_FDR_ = 0.003). The strongest negative effect between age and rs-FC was observed between areas in the occipital lobe: left lateral occipital cortex a – left lingual gyrus b (*ρ* = −0.37, *p*_FDR_ = 0.003), and left cuneus and pericalcarine cortex – left lateral occipital cortex a (*ρ* = −0.36, *p*_FDR_ = 0.003). The linear effect size was generally higher in the beta band compared to the alpha and theta frequency ranges.

In addition to the decreasing linear trend, a negative quadratic relationship was identified between participant age and beta-band rs-FC, most notably between central–parietal and central–posterior areas. The negative quadratic trend corresponds to an inverted U-shape, indicating that these connections strengthen in young adults through middle age and decline in older adults toward later life. This pattern resembles those observed in structural connectivity and fMRI functional connectivity networks (Betzel et al., 2014; Zhao et al., 2015). Some of the strongest quadratic associations were observed between left and right caudal middle frontal gyri 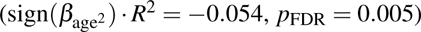, left supramarginal gyrus a and right inferior parietal cortex 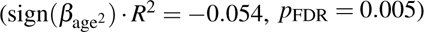, and left postcentral gyrus b and right lingual gyrus b 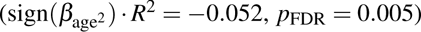.

In the gamma band, we observed increasing rs-FC trajectories as a function of age, without clear structure. The strongest effects were observed between the right middle temporal gyrus and the right lateraloccipital gyrus (*ρ* = 0.25, *p*_FDR_ = 0.004), as well as between the right superior temporal gyrus and the right lateral occipital gyrus (*ρ* = 0.23, *p*_FDR_ = 0.004). The gamma-band effects were generally weaker than effects in the other frequency bands. No significant quadratic associations between age and rs-FC were observed in the gamma band.

### Mean connectivity across the adult lifespan

To highlight age-related trends in rs-FC associated with individual brain regions, we calculated linear associations between the mean connectivity of each region and participant age in the delta, theta, alpha, beta, and gamma frequency bands (**Fig 3**).

**Figure 3.**
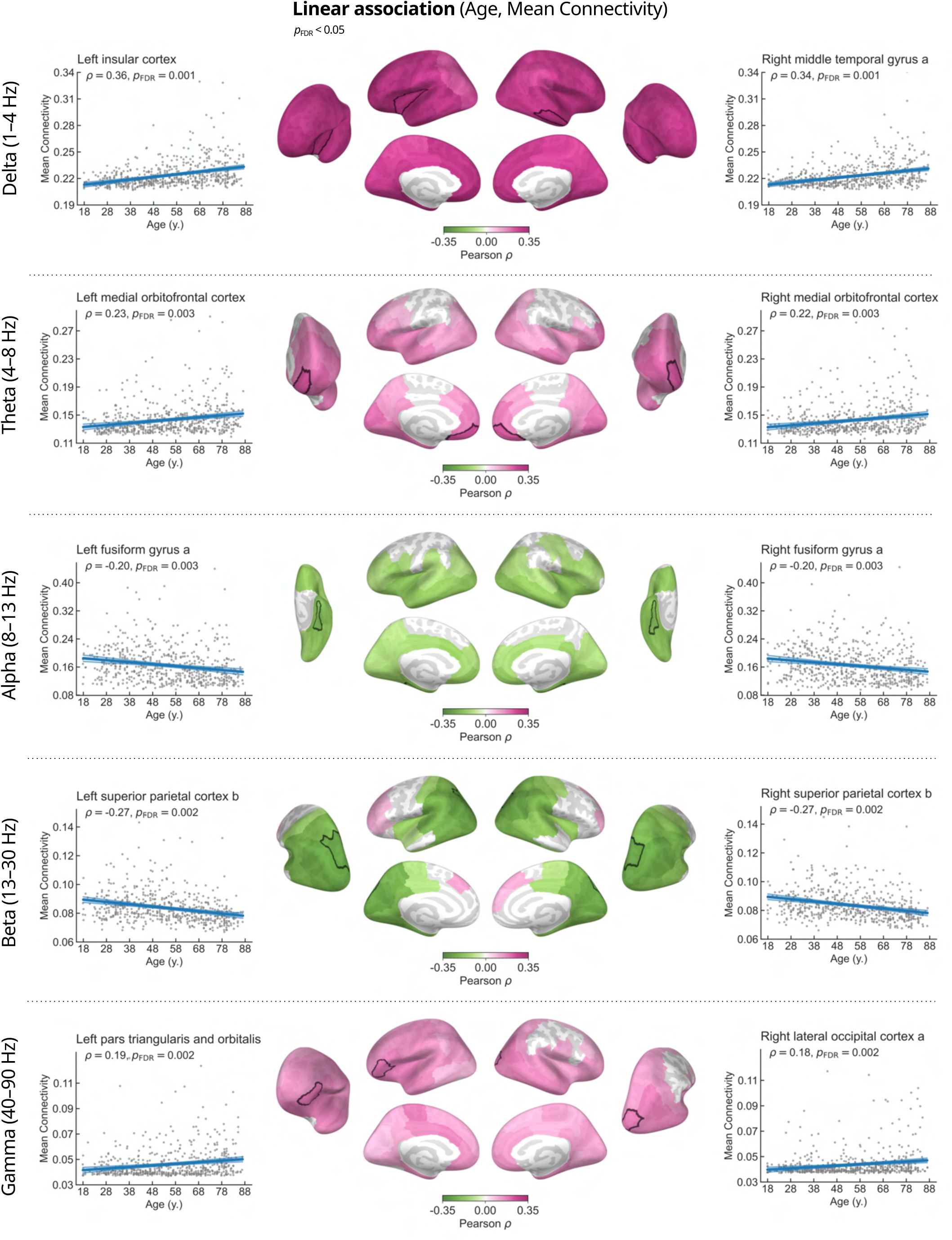
Mean connectivity changes across the adult lifespan. Brain regions showing significant associations between age and mean connectivity in the delta, theta, alpha, beta, and gamma frequency bands are colored according to their Pearson correlation coefficient. The cortical areas associated with the largest effect sizes in each frequency band are highlighted with white borders. The scatter plots in the left and right columns illustrate the linear relationships between participant age and mean connectivity of the highlighted regions. Each data point represents an individual subject. The lower intensity lines show the 95% confidence intervals.

In the delta band, statistically significant increases in mean connectivity were observed in all brain regions. The effects were strongest in the left insula (*ρ* = 0.36, *p*_FDR_ = 0.001) and right middle temporal gyrus (*ρ* = 0.34, *p*_FDR_ = 0.001). In the theta band, we observed a general increase in mean connectivity with age, particularly in frontal, temporal, and occipital areas. The strongest effect was observed bilaterally in the medial orbitofrontal cortices (left: *ρ* = 0.23, *p*_FDR_ = 0.003; right: *ρ* = 0.22, *p*_FDR_ = 0.003). In the alpha band, mean connectivity was negatively associated with age, and the effect was largest in the fusiform gyri (left: *ρ* = −0.20, *p*_FDR_ = 0.003; right: *ρ* = −0.20, *p*_FDR_ = 0.003). In the beta band, a more complex pattern showed age-related decreases in connectivity in the posterior and parietal regions combined with a modest positive trend in the frontal lobe. The largest effect size was associated with decreased connectivity of the bilateral superior parietal cortices (left: *ρ* = −0.27, *p*_FDR_ = 0.002; right: *ρ* = −0.27, *p*_FDR_ = 0.002). The gamma-band patterns were similar to those in the theta band, with positive linear trends in most brain regions, except parietal cortex. The strongest effects were observed in the left pars triangularis and orbitalis (*ρ* = 0.19, *p*_FDR_ = 0.002) and right lateral occipital cortex (*ρ* = 0.18, *p*_FDR_ = 0.002).

### Global connectivity and source power across the adult lifespan

To visualize and quantify age-related changes in whole-brain rs-FC across the 1–100-Hz frequency range, we calculated the linear and quadratic association spectra between participant age and global connectivity (**Fig 4**). We identified frequency-dependent patterns of both positive and negative linear and quadratic associations between age and global connectivity (**Fig 4B**). Generally, age and global connectivity were positively linearly associated at low frequencies below 7 Hz and high frequencies above 40 Hz. In contrast, a negative association was observed at intermediate frequencies (11–14 Hz and 20–23 Hz). The largest effect size was related to a negative association at 12 Hz (*ρ* = −0.32, *p*_FDR_ = 0.002), indicating reduced global connectivity with older age (**Fig 4C**). In addition to the linear trends across wide frequency ranges, we observed a negative quadratic association between age and global connectivity in a narrow 17–20-Hz range within the beta band, with the strongest effect at 17 Hz 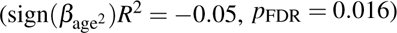.

**Figure 4.**
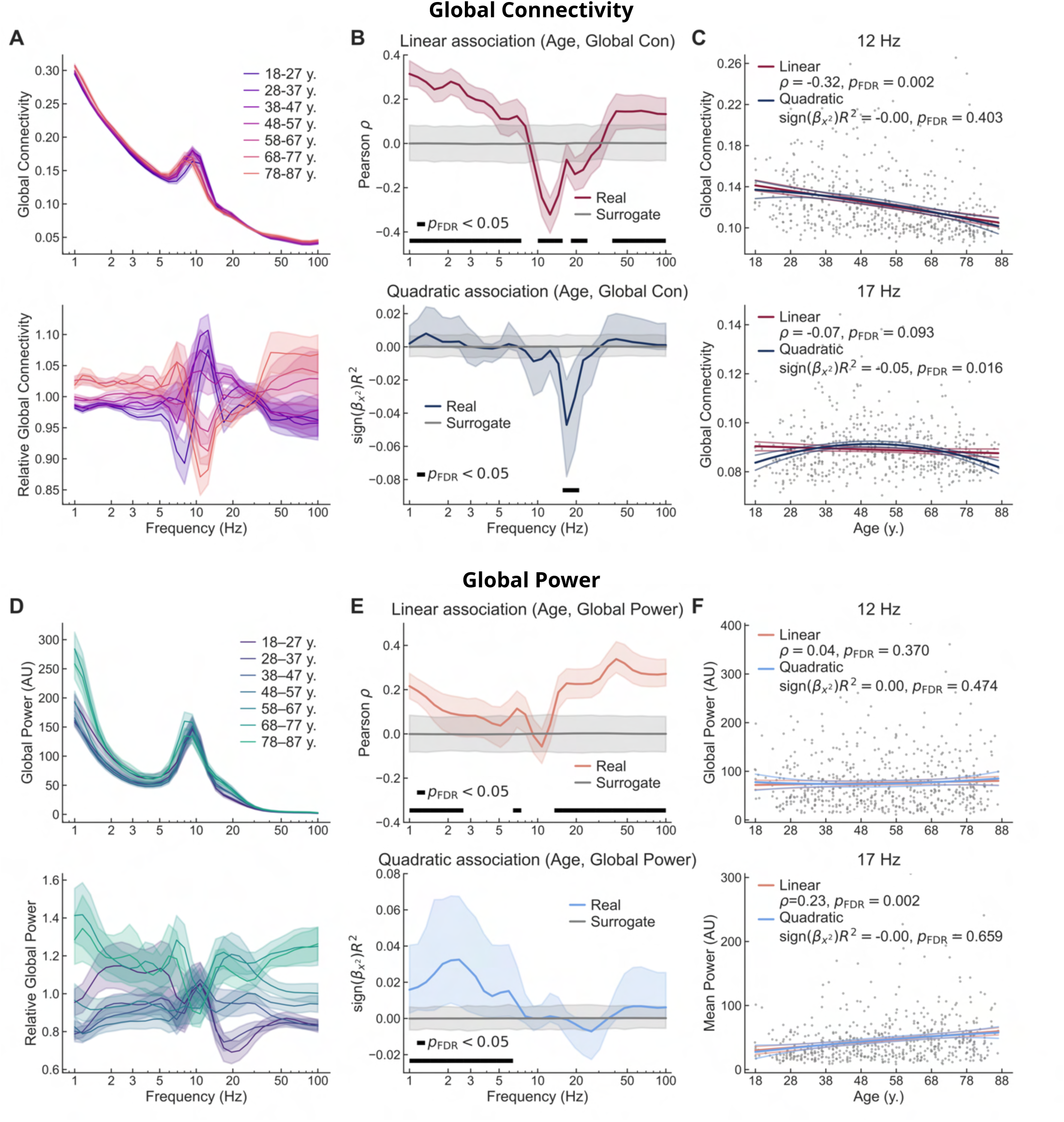
Global connectivity and source power change across the adult lifespan. (**A**) Group averages of absolute (top) and relative (bottom) global connectivity spectrum within age cohorts. The relative connectivity is normalized by the grand average in each frequency bin. Shaded areas indicate the standard error of the mean. (**B**) Linear (top) and quadratic (bottom) association spectra between participant age and global connectivity. Shaded areas indicate 95% bootstrap confidence intervals. (**C**) Scatter plots of global connectivity as a function of participant age at 12 Hz (top) and 17 Hz (bottom) show the peak linear and quadratic relationships, respectively. The lower intensity lines show the 95% confidence intervals. (**D**) Group averages of absolute (top) and relative (bottom) mean source power within age cohorts. The relative power is normalized by the grand average in each frequency bin. (**E**) Linear (top) and quadratic (bottom) association spectra between participant age and mean source power. (**F**) Scatter plots of mean source power as a function of participant age at 12 Hz (top) and 17 Hz (bottom) show non-significant or opposing trends compared to global connectivity.

To compare the adult lifespan trajectories of rs-FC to those of oscillatory power (i.e., local synchronization), we calculated the linear and quadratic association spectra between participant age and global source power (**Fig 4E**). We observed positive linear associations in low (1–2 Hz, 7 Hz) and high (14–100 Hz) frequencies. However, despite the associations between age and global connectivity, no significant linear relationships between age and global power were observed in the theta or alpha frequency ranges, except at 7 Hz. Positive quadratic associations between age and global source power were observed in low frequencies (1–6 Hz), again different from the frequency range where age and global connectivity were quadratically associated. This is illustrated in **Fig 4C** and **Fig 4F**, which highlight the absence of significant linear or quadratic associations between age and global source power at the frequencies of peak associations between age and global connectivity. These results demonstrate that rs-FC and oscillatory power contain non-redundant information and are not mere reflections of each other.

### Beta-band connectivity and sensorimotor attenuation

To investigate the relationship between rs-FC and sensorimotor attenuation, we calculated linear associations between the mean force overcompensation in the force matching task and the mean connectivity of each region in the beta band. We identified significant negative associations in the centroparietal regions comprising sensorimotor cortices, indicating that increased overcompensation, and thus greater sensorimotor attenuation, is related to lower connectivity in those areas (**Fig 5A**). In the left hemisphere, the strongest relationship was observed in the supramarginal gyrus (*ρ* = −0.21, *p*_FDR_ = 0.033). In the right hemisphere, the largest effect occurred in the postcentral gyrus (*ρ* = −0.20, *p*_FDR_ = 0.033). The distribution of mean force overcompensation contains outlier observations. To ensure that the results were not driven by outliers, we recomputed the analysis with the outliers excluded. Differences in the results were close to negligible (Supplementary Figure S4). Details are provided in Supplementary Material.

**Figure 5.**
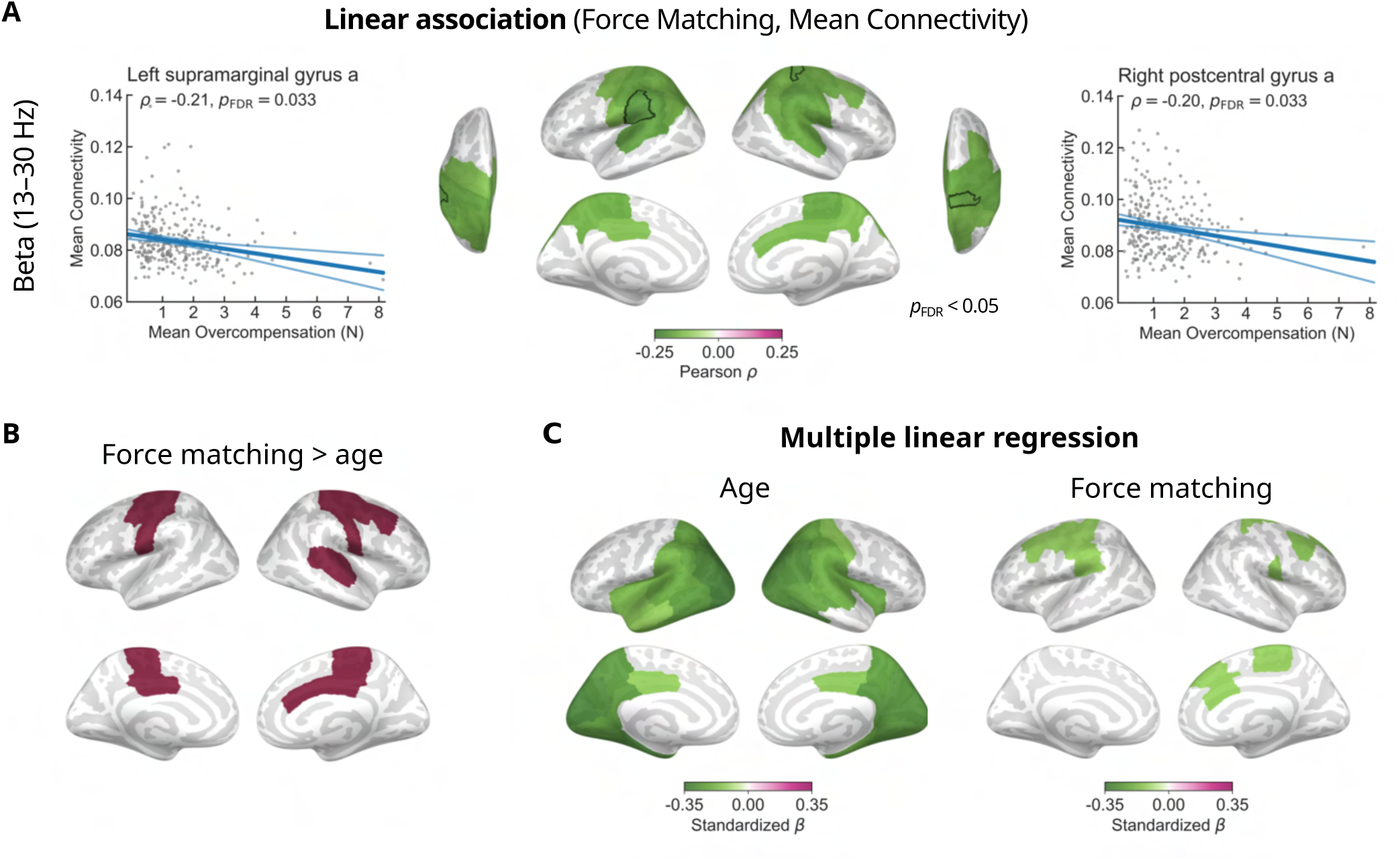
Association between mean connectivity and performance in the force matching task. (**A**) The brain regions that showed significant negative linear associations between beta-band mean connectivity and mean force overcompensation in the force matching task are depicted in green. The scatter plots present data from the regions with the strongest observed correlations—the left supramarginal gyrus and the right postcentral gyrus. The lower intensity lines show the 95% confidence intervals. (**B**) Brain regions where the linear association between beta-band mean connectivity and mean force overcompensation in the force matching task was over and above the linear association between beta-band mean connectivity and participant age are colored in red. (**C**) Effect sizes (standardized *β* -coefficients) of significant regions for age and mean overcompensation in the force matching task from a multiple linear regression model predicting mean connectivity.

To distinguish the relationship between rs-FC and sensorimotor attenuation from age-related effects, we identified the brain regions where the strength of the association between mean overcompensation in the force matching task and mean connectivity was over and above the strength of the association between participant age and mean connectivity (**Fig 5B**). This analysis identified clusters covering bilateral sensorimotor cortices, indicating that associations between rs-FC and sensorimotor attenuation are dissociable from the observed age-related decrease in beta-band rs-FC.

In addition to the mass univariate analysis, we conducted a standardized multiple linear regression analysis where both participant age and mean overcompensation in the force matching task were included as explanatory variables to predict mean connectivity in each region. **Fig 5C** shows the significant effect sizes with respect to both variables in each region. Similar to the univariate correlation-based analysis (**Fig 3** and **Fig 5A**), these results indicate that age-related effects are spatially distinct from sensorimotor effects. In particular, participant age affected mean connectivity mainly in posterior and temporal areas, while mean overcompensation in the force matching task was relevant in rolandic regions associated with sensorimotor control.

## Discussion

In this study, we aimed to elucidate the trends of resting-state functional connectivity (rs-FC) across the healthy adult lifespan and the potential role of rs-FC in sensorimotor integration. Our results revealed frequency-dependent rs-FC trajectories as a function of age. We also showed a linear relationship between the beta-band rs-FC and sensorimotor attenuation, indicating that the ongoing inter-areal synchronization in the beta band is involved in sensorimotor integration. Furthermore, we showed that rs-FC and source power have distinct frequency-dependent trajectories, suggesting that rs-FC provides additional information to that conveyed by source power alone.

In the beta band, our analysis of rs-FC network connections across the lifespan showed a linear age-related decrease in rs-FC between posterior and parietal regions. The observed effect size was larger in the beta band compared to the theta, alpha, and gamma frequency ranges. In addition to the linear trend, we observed an age-related negative quadratic ”inverted U-shape” trend between central and parietal regions, similar to what has been reported within resting-state networks in fMRI (Betzel et al., 2014), white matter volume (Bethlehem et al., 2022), and structural connectivity (Zhao et al., 2015). Future studies should investigate whether beta-band rs-FC reflects hemodynamic and structural coupling better than other frequency bands. Notably, linear and quadratic associations showed distinct spatial patterns, hinting that separate underlying mechanisms could drive these changes within the same frequency band. In general, the age-related patterns observed at the region level align with the results of a recent study by Stier and colleagues (2023) despite some methodological differences, increasing our confidence in the replicability of the results.

Beta-band oscillatory activity is central to sensorimotor control and has been associated with maintenance of the current sensorimotor state (Barone & Rossiter, 2021; Engel & Fries, 2010). Increased sensorimotor attenuation could be attributed to increased reliance on sensorimotor prediction as opposed to sensory feedback and has been associated with aging (Wolpe et al., 2016). Moreover, increased sensorimotor attenuation has been associated with increased fMRI functional connectivity between the supplementary motor area (pre-SMA) and the primary somatosensory cortex, along with decreased functional connectivity between the pre-SMA and frontostriatal areas (Wolpe et al., 2016). To investigate the underlying neuronal dynamics, we analyzed the relationship between rs-FC and mean overcompensation in the force matching task. In contrast to the fMRI findings, we observed an association between decreased connectivity of the broader sensorimotor areas in the beta band and increased force overcompensation. This association suggests that a higher level of beta-band functional connectivity at rest could be indicative of efficient sensory information processing, leading to less pronounced sensorimotor attenuation, whereas decreased beta-band connectivity could be related to increased sensory noise and greater emphasis on internal predictive models, observed as increased sensorimotor attenuation in healthy aging (Konczak et al., 2012; Moran et al., 2014). However, we also observed an age-related increase in beta-band rs-FC in the dorsomedial prefrontal cortex (**Fig 3**), which partly overlaps with the areas where increased force overcompensation was associated with decreased beta-band connectivity in the multiple linear regression analysis (**Fig 5C**). Since the dorsomedial prefrontal cortex has been associated with force control and motor planning (Coombes et al., 2012; Koechlin et al., 2000), this suggests that the neural circuits underlying increased force overcompensation in aging could be more complex than a mere decrease in rs-FC in sensorimotor areas. Furthermore, due to the cross-sectional nature of the current study, we were only able to show that rs-FC effects related to age and sensorimotor integration involve separate cortical areas. Future longitudinal work should further investigate whether age-related changes in functional connectivity in the beta band are causally linked to increased sensorimotor attenuation.

In the alpha band, our results showed an overall decrease in rs-FC with age, prominently between frontal, temporal, and occipital areas. At the region level, the strongest associations between age and mean connectivity were observed in the fusiform and parahippocampal gyri in the medial temporal lobe along with weaker effects in the wider frontal, temporal, and occipital areas. These results agree with a previous study that observed an age-related decrease in temporal lobe connectivity in the same cohort (Stier et al., 2023). Reduced alpha-band rs-FC has previously been associated with reduced VSTM precision in healthy aging (Sahoo et al., 2020) and is widely considered a marker of mild cognitive impairment and Alzheimer’s disease (AD)(Buzi et al., 2023; López et al., 2014; Mandal et al., 2018; Stam et al., 2006). Although one could draw parallels between the patterns observed in healthy aging and neuropathologies such as AD, it has also been reported that the evolving rs-FC patterns differ between healthy aging and AD (Koelewijn et al., 2017). However, in the absence of direct mechanistic evidence, it remains challenging to interpret the neurophysiological mechanisms underlying the changing patterns in aging. To that end, neuronal dynamics producing realistic rs-FC estimates in the alpha band have been simulated using neural mass models (David et al., 2004). Several studies combining diffusion tractography data and neural mass models of varying complexity have shown that structural connectivity predicts functional connectivity, but only to a limited degree (Abeysuriya et al., 2018; Forrester et al., 2024; Griffiths et al., 2020; Kulik et al., 2023; Pons et al., 2010; Ponten et al., 2010). Structural reorganization of the brain, particularly that of thalamocortical connections, could thus partly explain the observed trajectories. Furthermore, similarity between structural connectivity and empirical alpha-band rs-FC has been linked to cognitive performance in the elderly (Jauny et al., 2024). Nevertheless, it is unlikely that structural connectivity alone would explain all observed functional changes.

It is well known that the frequency of the individual alpha peak decreases throughout the adult lifespan from approximately 10 Hz in young adulthood to less than 9 Hz in the elderly (Klimesch, 1999; Scally et al., 2018), although considerable inter-individual variability is involved (Haegens et al., 2014). The decrease of the peak alpha frequency is a possible confounding factor in functional connectivity analysis, as the strongest connectivity is typically observed at the peak frequency (Scally et al., 2018). Age-related changes in source power could also lead to biased estimates of age-related changes in connectivity since higher source power translates to a higher signal-to-noise ratio (SNR), and it is known that variations in SNR are a major source of bias in functional connectivity analysis (Bastos & Schoffelen, 2016; Donoghue et al., 2022). Although prior work suggests age-related increases in MEG source power in the alpha band (Rempe et al., 2023; Stier et al., 2023), we did not observe age-related trends in global power in that frequency range despite an observed decrease in global connectivity. These results together suggest that our rs-FC findings in the alpha band are not merely echoes of altered source power. Similarly, we observed no significant linear changes in theta power with age, and only a linear increase in beta power despite both negative linear and negative quadratic trends in rs-FC. Moreover, while global power in the delta band showed a positive quadratic trend, only a linear trend was observed in delta rs-FC. Thus, we provide evidence that global trajectories of inter-areal functional connectivity are distinct from those of local synchronization (i.e., spectral power).

In our analysis of age-related changes in all-to-all connectivity, we observed increased rs-FC in the low-frequency delta and theta bands in older age, particularly between frontal–temporal and frontal–occipital areas, with a notable contribution of interhemispheric coupling. This increase was mirrored by the region-level mean connectivity results, which showed increased delta-band connectivity in all cortical areas, and increased theta-band connectivity in frontal, temporal, and occipital areas, with the largest effects in the bilateral medial orbitofrontal cortices. These findings in the theta band are in line with a previous study by Stier and colleagues (2023), although our results suggest a more pronounced role for the frontal cortex. The functional significance of these age-related changes in spontaneous low-frequency functional connectivity remains unclear, since most studies have either been task-based or compared rs-FC in healthy aging to that in neurodegenerative disorders. Theta-band connectivity, particularly that of the medial prefrontal cortex, has been credited with a central mediating role in cognitive control and working memory through the entrainment of distal regions (Cavanagh & Frank, 2014; Daume et al., 2017; Liebe et al., 2012). While these effects were observed during task performance, there is also evidence that the level of theta rs-FC could reflect the functioning of attentional mechanisms (Fellrath et al., 2016). Moreover, it has been suggested that increased delta–theta functional connectivity could have a compensatory role in cognitive aging, allowing for maintenance of cognitive performance in older age (Jauny et al., 2022b).

In the gamma band, we observed increased rs-FC with older age. The effects were generally weaker than in the other frequency bands and did not show clear spatial patterns in the all-to-all connectivity analysis (Fig 2). However, although previous studies have also observed age-related increases in long-range gamma-band rs-FC (Doval et al., 2024; Stier et al., 2023), we advise caution when interpreting these results. In fact, early modeling work has suggested that gamma oscillations occur predominantly in local neuronal circuits and cannot support long-range connections involving long conduction delays (Kopell et al., 2000). More recently, other studies have observed long-range functional connectivity in the gamma band, e.g., increased long-range gamma-band coherence between local field potential recordings of visual areas V1 and V4, as well as the frontal eye field and V4, during visual attention tasks (Bosman et al., 2012; Gregoriou et al., 2009; Grothe et al., 2012), and task-dependent modulation of gamma-band coherence between the spinal cord and the motor cortex (Schoffelen et al., 2005). These investigations of long-range gamma-band functional connectivity have been limited to specific sensory and motor circuits, where they could represent a bottom-up attentional mechanism (Bastos et al., 2015; Fries, 2015). On the contrary, evidence for a functional role of spontaneous long-range gamma synchronization is lacking. Hence, the underpinnings and consequences of increased gamma rs-FC in older age remain elusive.

Although we addressed most of the methodological concerns inherent to estimating resting-state functional connectivity, one should be aware of a few remaining limitations when interpreting the results. First, many functional connectivity metrics exist, and they often produce divergent results. We repeated our rs-FC analysis using amplitude envelope correlation (AEC) and found both similarities and differences between AEC and WPLI results (Supplementary Material). Therefore, caution should be exercised when comparing results across studies that utilize different metrics. Moreover, although the applied connectivity metric, Weighted Phase-Lag Index (WPLI), is insensitive to instantaneous synchronization and resilient against noise (Vinck et al., 2011), spurious interactions between sources near the truly interacting sources could remain in the results (Palva et al., 2018). This limitation should be considered when interpreting the observed spatial patterns. Furthermore, WPLI emphasizes phase differences near 90 and 270 degrees to reduce the contributions of putatively noisy near-zero phase differences. However, while FC estimators should consider neuronal conduction delays and thus non-zero phase lags (Fries, 2015), the specific preference for 90- and 270-degree phase differences is not strictly motivated biologically. Another potential confounding factor arises from the uncontrolled nature of resting-state measurements; it is not well understood how the internal cognitive state and vigilance of the participant affect rs-FC, although it has been suggested that drowsiness does not significantly affect phase-based estimates of functional connectivity (Strijbis et al., 2022). Finally, the large interindividual variability observed in our study could limit the interpretation of rs-FC analysis at the individual level. Nevertheless, the use of large MEG datasets enables modeling of the normal healthy variability in functional connectivity, which could lead to improved diagnostics and prognosis of brain disorders (Itälinna et al., 2023). Our demonstration of age-related variability in rs-FC emphasizes the importance of age-stratified normative modeling.

An additional limitation of our study is that we do not have information on potential asymptomatic neuropathological changes, such as amyloid-beta or tau protein deposition, in the elderly cohort. While considered a characteristic marker of clinical Alzheimer’s disease (AD) (LaFerla et al., 2007), amyloid-beta deposition is observed in 10–30% of asymptomatic healthy elderly (Chételat et al., 2013). Through various cellular mechanisms, amyloid-beta deposition alters the excitation/inhibition (E/I) balance towards increased cortical excitability, which could compromise large-scale neuronal networks (Busche & Konnerth, 2016; Maestú et al., 2021). Previous studies have associated increased delta and theta-band and decreased alpha-band rs-FC in MEG with amyloid-beta deposition in both clinical AD patients (Schoonhoven et al., 2022) and healthy elderly (Nakamura et al., 2017). Others have reported an absence of significant associations between amyloid accumulation and EEG rs-FC in healthy elderly (Teipel et al., 2018). Although the individuals in our cohort were cognitively normal (MMSE score ¿ 24, no neurological diagnoses), we cannot rule out a potential contribution of preclinical amyloid-beta deposition in the elderly group, since it can start decades before clinical symptoms appear (Jansen et al., 2015) and the functional changes previously associated with amyloid-beta deposition overlap with our age-related findings. Future studies on age-related effects in rs-FC should therefore consider controlling for amyloid deposition. Our study contributes to the literature on healthy aging with three main findings. First, we showed that resting-state functional connectivity (rs-FC), quantified as phase coupling between spontaneous cortical oscillations, is altered in healthy aging. The observed changes showed different spatial patterns in the delta, theta, alpha, beta, and gamma frequency bands. In the beta band, rs-FC followed an inverted U-shape trajectory with age, mirroring patterns previously observed in MRI studies of structural and functional connectivity. Second, we showed that the global trends in rs-FC differ from those of oscillatory source power, alleviating concerns that the age-related changes in rs-FC would be confounded by changes in source power. Finally, our results indicate an association between reduced beta-band rs-FC in sensorimotor areas and increased sensorimotor attenuation, which suggests that rs-FC could play a role in the integration of sensory signals and internal predictive models. In summary, our results contribute to the understanding of the role of brain oscillations in healthy aging and open new avenues for future studies linking the observed changes in rs-FC with behavior and cognition, as well as further elucidating the putative role of beta-band rs-FC in sensorimotor integration.

## Data and code availability

The original MEG, MRI, and behavioral data are available upon request from the Cam-CAN data repository (https://camcan-archive.mrc-cbu.cam.ac.uk//dataaccess/). The code developed and used in this study is available upon request from the corresponding author.

## Acknowledgments

Data collection and sharing for this project were provided by the Cambridge Center for Ageing and Neuroscience (Cam-CAN). Cam-CAN funding was provided by the UK Biotechnology and Biological Sciences Research Council (grant number BB/H008217/1), together with support from the UK Medical Research Council and University of Cambridge, UK. S.R. and L.P. disclose support for the research of this work by the National Institute for Neurological Disorders and Stroke (grant number R01NS104585), and the Helsinki–Uusimaa Regional Council. J.C.A.-D. and L.P. disclose support for the research of this work by Business Finland (grant “DIGIMIND” 7981/31/2022) and the Norman Loveless Memorial Fund. M.L. discloses support for the research of this work by the Swedish Cultural Foundation in Finland. We thank Matti S. Hämäläinen for providing comments on the manuscript and acknowledge the computational resources provided by the Aalto Science-IT project.

## Ethics approval statement

The primary study responsible for data collection was conducted in accordance with the Declaration of Helsinki and was approved by the Cambridgeshire 2 Research Ethics Committee (reference: 10/H0308/50). The participants gave written informed consent for the study. These data were shared with the authors by the Cambridge Center for Ageing and Neuroscience (Cam-CAN) and were handled in accordance with the Cam-CAN data use agreement.

