## Supplementary Material for "Frequency-resolved cortical functional connectivity across the adult lifespan"

### Similarities and differences between amplitude envelope correlation and WPLI

The study of cortical functional connectivity using non-invasive electrophysiological measurement techniques, MEG and EEG, is not straightforward due to intrinsic properties of these signals, namely poor signal-to-noise ratio (SNR) and linear mixing between signals from different source neuronal populations. Interactions between brain regions cannot be directly measured, but rely on statistical measures of similarity between the signals. This has led to the development of a spectrum of functional connectivity metrics (Bastos & Schoffelen, 2016; Wang et al., 2014). There is currently a large variability in results across studies utilizing different metrics (Jensen et al., 2025) and substantial differences in test–retest reliability across metrics (Colclough et al., 2016; Garcés et al., 2016; Marquetand et al., 2019).

The functional connectivity metric applied in the current study, the weighted phase-lag index (WPLI), is a measure of phase synchronization between frequency-resolved signals. However, numerous studies on rs-FC have also applied power or amplitude envelope correlations (AEC) between band-restricted signals (Brookes et al., 2011; Colclough et al., 2015; Coquelet et al., 2017; Hipp et al., 2012; Khan et al., 2018; Schoonhoven et al., 2022). To facilitate comparison with previous and future AEC studies, we repeated our rs-FC analysis using AEC on the same narrowband analytic signals used to estimate WPLI. We calculated AEC between pairwise-orthogonalized amplitude envelopes to avoid artificial zero-lag interactions (Hipp et al., 2012).

The all-to-all AEC results are shown in Supplementary Figure S1. In the delta band, the results were similar to WPLI, with age-related linear increases of functional connectivity strongest between frontal, temporal, and occipital regions. In the theta band, increased frontal–temporal rs-FC in older age was also observed, but unlike with WPLI, we did not observe increased frontal–occipital coupling. Instead, weak linear decreases in rs-FC between occipital areas were found. In the alpha and beta bands, the identified linear patterns largely differed from those observed using WPLI. Although decreased rs-FC in older age between posterior regions was found using both AEC and WPLI, AEC showed linear increases in rs-FC between the prefrontal cortex and other regions. Nevertheless, both AEC and WPLI indicated inverted U-shaped quadratic trajectories in the beta band. In the gamma band, no significant associations between age and AEC were observed.

In addition to all-to-all connectivity, we found differences and similarities between WPLI and AEC at the region (Supplementary Figure S2) and global (Supplementary Figure S3) levels. In particular, fewer significant associations were observed in the AEC analysis. While all brain regions showed increased delta-band AEC, no significant relations between theta or gamma AEC and age were found. Moreover, in the alpha band, only the dorsomedial prefrontal cortex, and in the beta band, only prefrontal areas and the left middle temporal gyrus showed significant positive linear associations with participant age. These results contrast the WPLI findings, where significant reductions in alpha and beta rs-FC were indicated in large parts of the cortex. At the global level, both AEC and WPLI increased linearly in the delta range. However, besides the delta band, the only other significant age-related linear trajectory for global AEC was an increasing one at 14 Hz, in contrast to downward trajectories found in the alpha and beta ranges for WPLI. Nevertheless, both AEC and WPLI indicated inverted U-shaped global trends in the beta band.

The observed differences highlight the divergent sensitivity profiles of AEC and WPLI, which could capture non-redundant information and different underlying neuronal processes (Siems & Siegel, 2020). This was not surprising, as previous studies have also reported contradictory results between amplitude correlations and phase synchronization measures (Schoonhoven et al., 2022). Our results suggest that AEC might be less sensitive to age-related changes than WPLI. A previous study by Coquelet and colleagues (2017) with a smaller sample size found no changes in power envelope correlations in healthy aging, which could be a result of this lower sensitivity. On the other hand, AEC could have higher test–retest reliability (Colclough et al., 2016), which is crucial for biomarker applications. To best describe age-related and pathological changes in rs-FC, it can be beneficial to apply several measures and thereby leverage their complementary qualities.

### Outliers in the force matching task

The distribution of mean force overcompensation contains outlying observations (main Fig **??**). To ensure that the results were not driven by outliers, we recomputed the analysis with the outliers excluded. To that end, we recomputed the analysis with the data points outside [Q1–1.5IQR, Q3+1.5IQR] removed, where Q1 and Q3 are the first and third quartiles, and IQR is the interquartile range. Since the number of included data points was still large (277 data points vs. 284 data points before the exclusion), the corresponding differences in the results were close to negligible. While 30 out of 90 regions were significant before the outliers were excluded, 28 remained afterwards. Specifically, the regions left posterior cingulate cortex, right caudal anterior cingulate cortex, right inferior parietal cortex a, and right superior parietal cortex b were not significant after outliers were removed, while the regions left superior temporal gyrus a and left middle temporal gyrus a were elevated to significance. Supplementary Figure S4 shows the results with the outliers excluded.

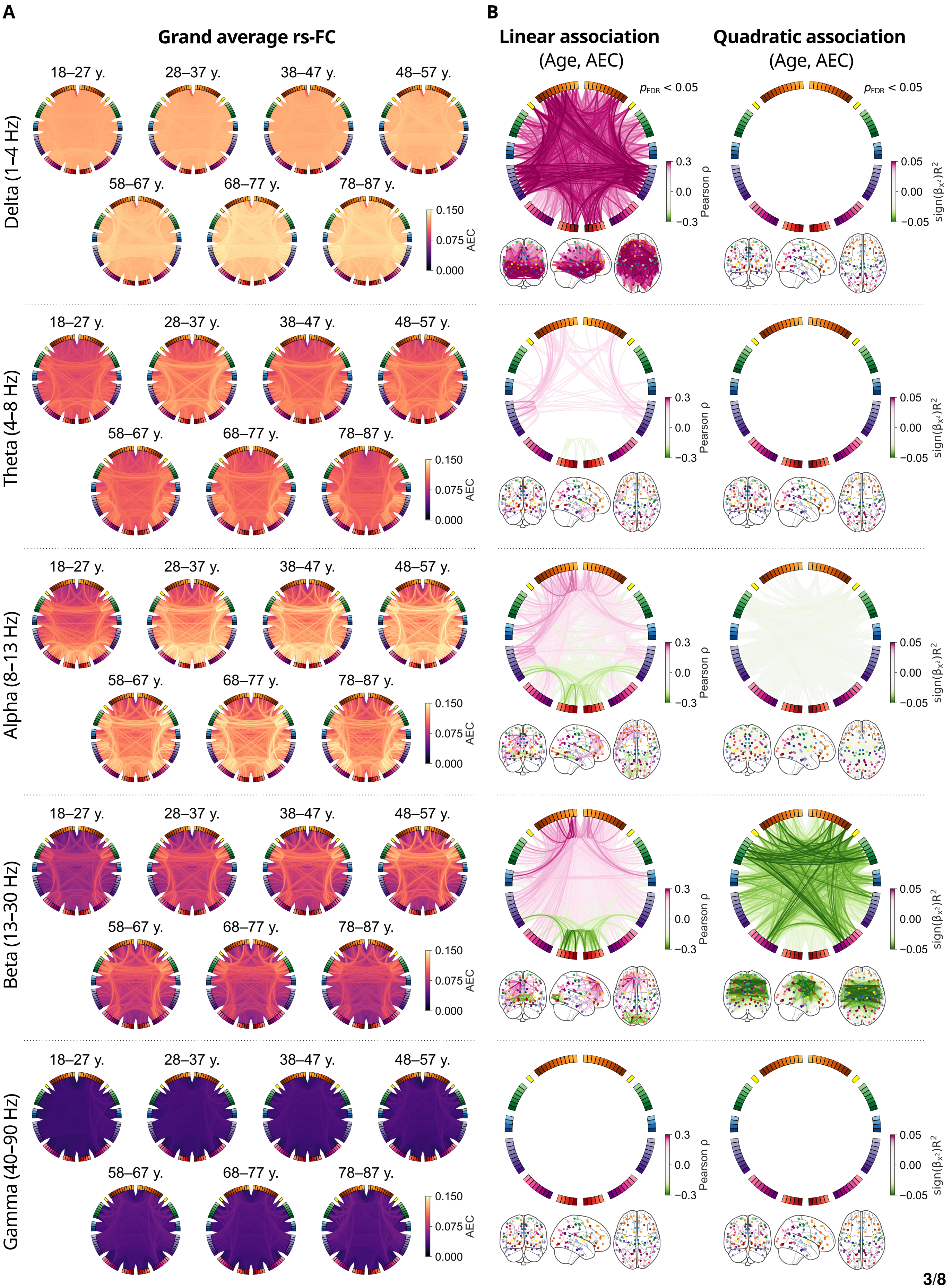

A
Grand average rs-FC
18–27 y.
28–37 y.
38–47 y.
48–57 y.
58–67 y.
68–77 y.
78–87 y.
Delta (1–4 Hz)
Theta (4–8 Hz)
Alpha (8–13 Hz)
Beta (13–30 Hz)
Gamma (40–90 Hz)
0.150
0.075
0.000
AEC
B
Linear association
(Age, AEC)
Quadratic association
(Age, AEC)
$p_{FDR} < 0.05$
0.3
0.0
−0.3
Pearson ρ
0.05
0.00
−0.05
$sign(\beta_{x^2})R^2$

**Supplementary Figure S1. Amplitude envelope correlation (AEC) connectivity changes across the adult lifespan.** (**A**) Group averages of AEC functional connectivity within age cohorts in the delta (1–4 Hz), theta (4–8 Hz), alpha (8–13 Hz), beta (13–30 Hz), and gamma (40–90 Hz) frequency bands show distinct frequency-dependent patterns. Each circular plot shows all links of the unthresholded connectivity graph. (**B**) Linear (left column) and quadratic (right column) associations between participant age and connection strength reveal frequency-band-specific patterns of change across the adult lifespan. Each colored line in the figure represents the effect size of a significant association; the Pearson correlation coefficient for linear associations, and the coefficient of determination multiplied by the sign of the quadratic term coefficient for the quadratic associations. The circular plots and schematic brains provide complementary visual perspectives of the same data.

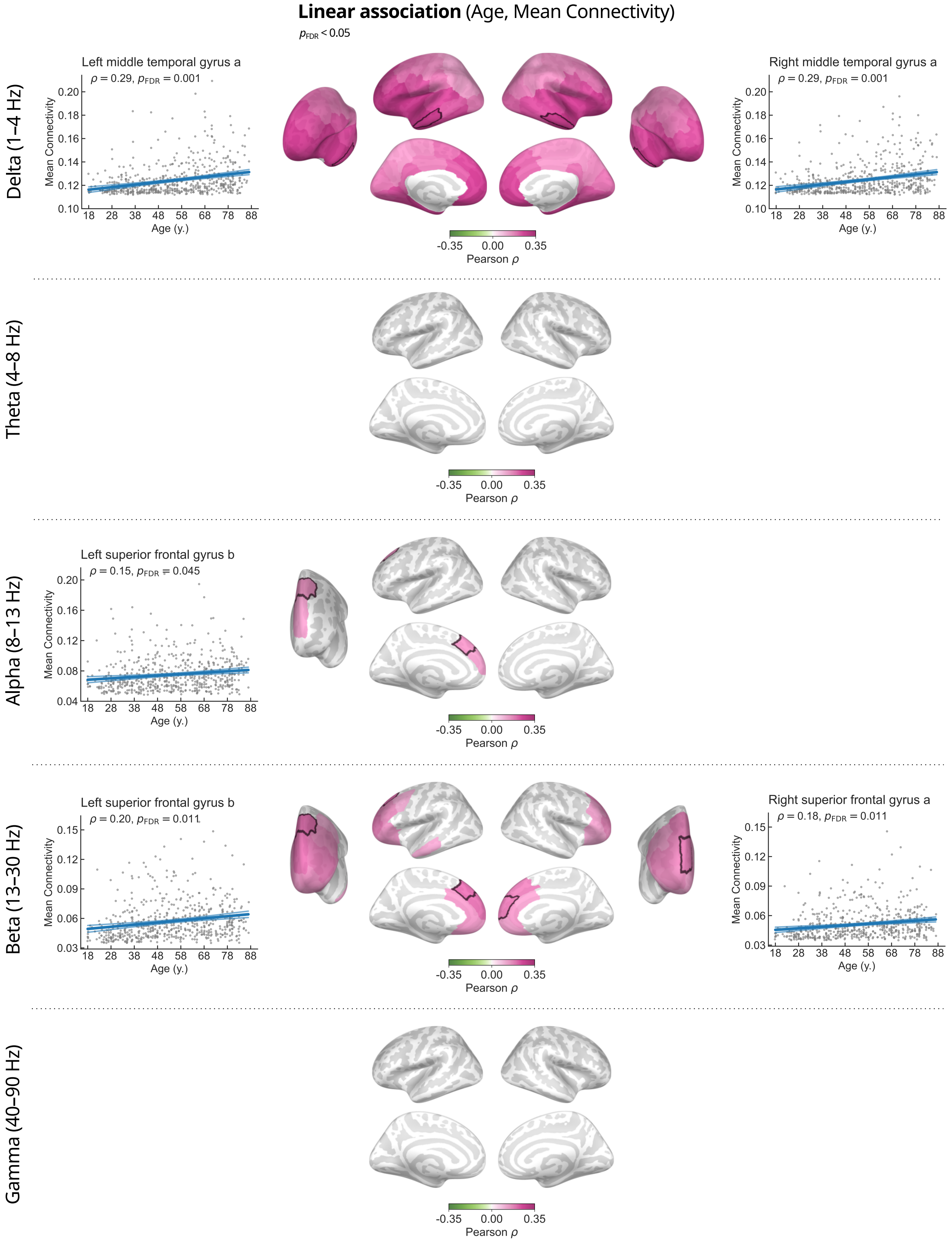
Linear association (Age, Mean Connectivity)
$p_{FDR} < 0.05$
Delta (1–4 Hz)
Left middle temporal gyrus a
$\rho = 0.29$, $p_{FDR} = 0.001$
Right middle temporal gyrus a
$\rho = 0.29$, $p_{FDR} = 0.001$
Theta (4–8 Hz)
Alpha (8–13 Hz)
Left superior frontal gyrus b
$\rho = 0.15$, $p_{FDR} = 0.045$
Beta (13–30 Hz)
Left superior frontal gyrus b
$\rho = 0.20$, $p_{FDR} = 0.011$
Right superior frontal gyrus a
$\rho = 0.18$, $p_{FDR} = 0.011$
Gamma (40–90 Hz)
Mean Connectivity
Age (y.)
18 28 38 48 58 68 78 88
-0.35 0.00 0.35
Pearson $\rho$

**Supplementary Figure S2.** **Mean amplitude envelope correlation (AEC) connectivity changes across the adult lifespan.** Brain regions showing significant associations between age and mean connectivity in the delta, alpha, and beta frequency bands are colored according to their Pearson correlation coefficients. The cortical areas associated with the largest effect sizes in each frequency band are highlighted with white borders. The scatter plots in the left and right columns illustrate the linear relationships between participant age and mean connectivity of the highlighted regions. Each data point represents an individual subject. The lower intensity lines show the 95% confidence intervals.

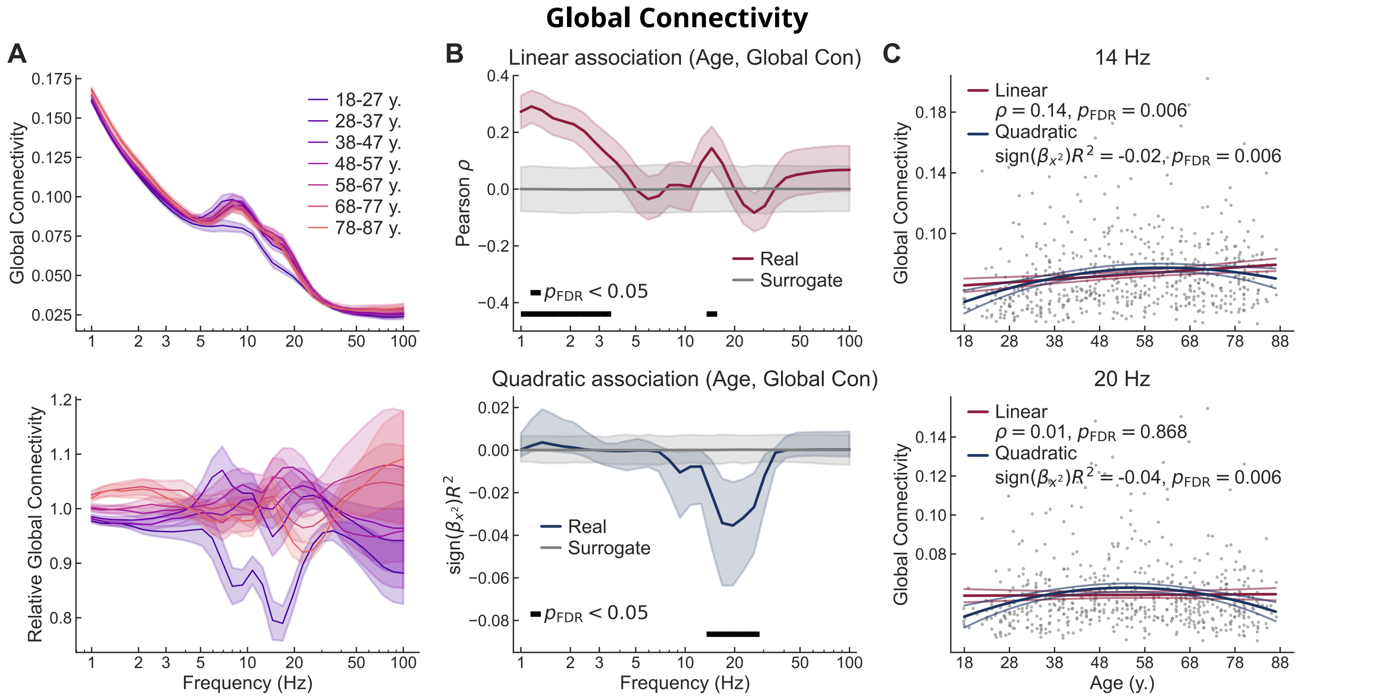


**Supplementary Figure S3.** **Global amplitude envelope correlation (AEC) changes across the adult lifespan.** (**A**) Group averages of absolute (top) and relative (bottom) global AEC connectivity spectrum within age cohorts. The relative connectivity is normalized by the grand average in each frequency bin. Shaded areas indicate the standard error of the mean. (**B**) Linear (top) and quadratic (bottom) association spectra between participant age and global connectivity. Shaded areas indicate 95% bootstrap confidence intervals. (**C**) Scatter plots of global connectivity as a function of participant age at 12 Hz (top) and 17 Hz (bottom) show the peak linear and quadratic relationships, respectively. The lower intensity lines show the 95% confidence intervals.

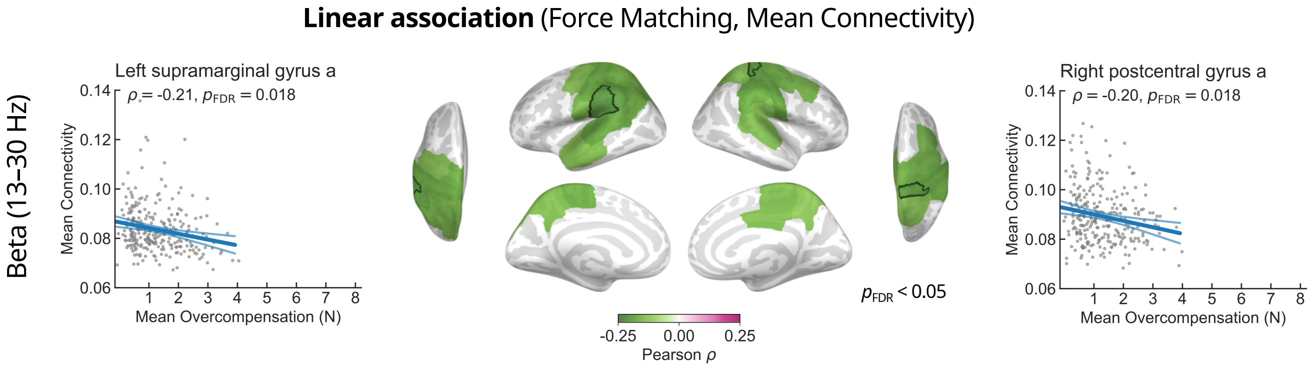


**Supplementary Figure S4. Association between mean connectivity and performance in the force matching task with outliers excluded.** (**A**) The brain regions that showed significant negative linear associations between beta-band mean connectivity and mean force overcompensation in the force matching task are depicted in green. Participants with mean overcompensation scores outside [Q1-1.5IQR, Q3+1.5IQR] have been excluded. The scatter plots present data from the regions with the strongest observed correlations—the left supramarginal gyrus and the right postcentral gyrus. The lower intensity lines show the 95% confidence intervals.
